# Aligning Bacteria and Synthetic Biomolecules with Engineered DNA Fibers

**DOI:** 10.1101/2020.12.28.423604

**Authors:** Jonathan R. Burns

## Abstract

DNA nanotechnology enables user-defined structures to be built with unrivalled control. However, the approach is currently restricted across the nanoscale, yet the ability to generate macroscopic DNA structures has enormous potential with applications spanning material, physical and biological science. I have employed DNA nanotechnology^[1, 2]^ and developed a new macromolecular nanoarchitectonic^[3]^ assembly method to produce DNA fibers with customizable properties. The process involves coalescing DNA nanotubes under high salt conditions to yield filament superstructures. Using this strategy, fibers over 100 microns long, with stiffnesses 10 times greater than cytoskeletal actin filaments can be fabricated. The DNA framework enables fibers to be functionalized with advanced synthetic molecules, including, aptamers, origami, nanoparticles and vesicles. In addition, the fibers can act as bacterial extracellular scaffolds and align *E.coli* cells in a controllable fashion. The results showcase the opportunities offered from DNA nanotechnology across the macroscopic scale. The new biophysical approach should find widespread use, from the generation of hybrid-fabric materials, platforms to study cell-cell interactions, to smart analytical and purification devices in biomedicine.

## 1.0 Introduction

DNA nanotechnology has produced significant advancements across materials science.^[1]^ The DNA origami construction approach enables discrete structures with nanometer control to be developed readily.^[2]^ The applications are widespread and include robots with programmable movement,^[4]^ DNA-based nanopores with charge selective ion transport,^[5]^ to smart sensing devices which deliver cargo to designated sites *in vivo*.^[6, 7]^ More recently, the field has been expanded towards supramolecular structures which assemble across the low micron scale.^[8]^ Using recognition sites, DNA origami blocks are able to combine into spherical superstructures,^[9]^ organized frameworks can shape lipid vesicles^[10]^ and plate arrays which assemble on 2D surfaces to mimic famous artwork.^[11]^ This emerging field is opening the way towards novel nanoarchitectonic structures^[3]^ with unique nanoscale functionality. However, expanding this concept further across the micron scale will enable new macromolecular materials to be assembled with unsurpassed biophysical control. These materials may include engineered woven fabrics with tailored nano-properties, synthetic actin filaments that simulate cellular movement inside model cells, to extracellular scaffolds which organize cells to aid biomedical research.^[12]^

DNA-based nanotubes are well suited as nanoarchitectonic structures. DNA strands can be programmed to form extended 1D lattices. To date, a diverse array of nanotubes have been engineered with controllable diameters and inter-duplex connectivity.^[13, 14, 15–17]^ The arrays can be functionalized and support the 1D organization of metal ions,^[18]^ plasmonic particles,^[19]^ quantum dots^[20]^ and fluorescent dyes with tailored properties.^[21]^ In addition, DNA nanotubes can be assembled into higher order nanofibers using specially designed junctions or organic modifications.^[17, 22]^ However, DNA nanotubes and nanofibers are highly flexible across the micron scale which limits their macroscopic potential. Therefore an alternative strategy is required to rigidify DNA nanotubes.

DNA condensation is a natural phenomenon. In order for genomic DNA to fit inside the nucleus of a cell, histone complexes to DNA to form a condensed superstructure.^[23]^ This phenomenon has been exploited in materials science, for example, Bar-Ziv and co-workers generated 1D patterned bio-chips using spermidine-DNA duplex condensates.^[24]^ Inspired by the favorable properties of condensation, I have developed a novel macromolecular assembly method which condenses flexible DNA nanotubes into rigid fibers (Figure 1). High concentrations of divalent metal cations complex between adjacent phosphate anions to form a stable inter-tube complex (Figure 1a, inset). The component nanotube was first presented by Winfree and co-workers^[13]^ and is assembled from subunits comprised of 5 oligonucleotides. Complementary toe-hold single strands positioned at each corner of the subunit overlap with a pitch of 25.8 ° to form a 7 subunit, 14 helix bundle. The bundles elongate upon annealing to generate a flexible nanotube framework (Figure S1-3).

**Figure 1.**
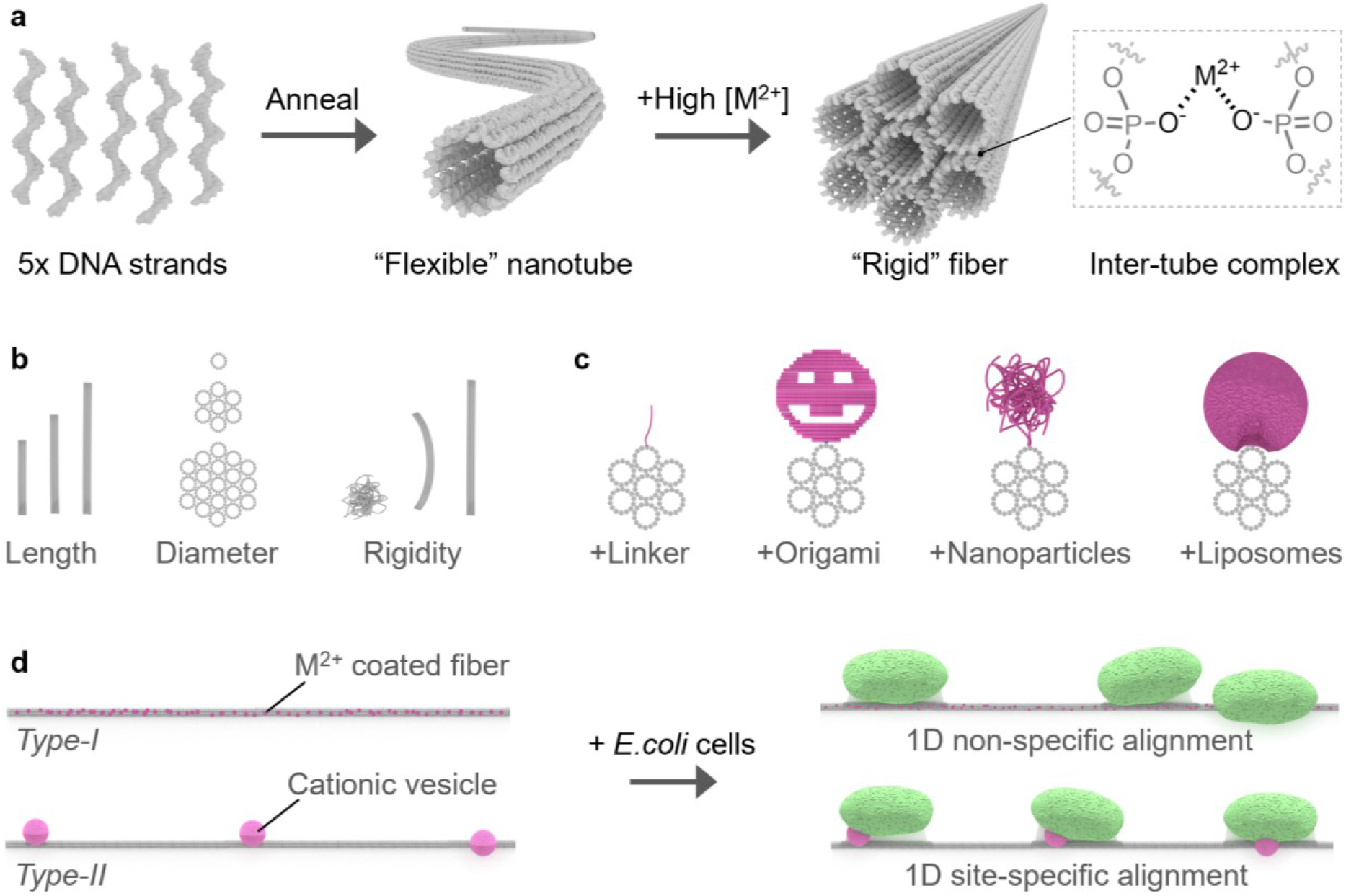
Overview of engineered DNA fibers. **a)** 5 component oligonucleotides anneal to form flexible DNA nanotubes under low salt conditions, or highly rigid DNA fibers (grey) in the presence of high concentrations of divalent metal cations, chemical structure of fiber inter-nanotube metal complex (inset). **b)** Fibers are highly versatile and can be customized in length, diameter and rigidity. **c)** Fibers can be decorated with advanced synthetic molecules, including, DNA linkers, DNA origami, DNA nanoparticles and liposomes (magenta). **d)** DNA fibers can align *E.coli* cells (green) *via* type-I and type-II mechanisms, generating non-site-specific and site-specific (magenta spheres) 1D extracellular frameworks, respectively.

Structural analysis of the nanotubes and fibers was conducted first to identify their core parameters, including diameter, length and rigidity (figure 1b). The superstructures’ metal cation concentration dependency was established, along with annealing profiles, optimal annealing rates and biocompatibility. Next, their modular properties were studied, and how the introduction of π-stacking cyanine dyes, or multiple seed-annealing cycles influences fiber formation. Following this, the fiber assembly process was applied to different DNA nanotube designs to confirm the methodology is generalizable. To showcase the DNA-enabled approach, a range of advanced synthetic biomolecules were incorporated (Figure 1c). The engineered filaments were then used to bind bacteria cells to function as extracellular scaffolds (Figure 1d). Two cell aligning mechanisms were investigated, a non-site-specific alignment (type-I) using divalent metals cations, and a site-specific alignment (type-II) using cationic liposomes. Finally, advanced bacterial cell matrix applications were developed, including alignment under complex multicellular conditions to organized cell division along the framework.

## 2.0 Results

### 2.1 Structural characterization of DNA nanotubes and fibers

The DNA framework must be sufficiently stable and rigid across the micron range for macroscopic applications. To achieve this, individual DNA nanotubes can be coalesced into structurally more rigid and longer fibers. Experimental characterization shows isolated DNA nanotubes form readily after mixing the 5 component strands and annealing less than 1 hr in buffer containing 14 mM MgCl2 (see Figure S1-3, S7-12, Table S1-3). Transmission electron microscopy (TEM) and confocal laser scanning microscopy (CLSM) confirmed the DNA nanotube formation (Figure 2a, S13-14). Nanotubes with diameters of 18.3 nm ± 2.2 nm are observed, which is comparable to previously published results.^[13]^ However, the frameworks are highly flexible and form large interwoven bundles measuring ~ 6 μm across (Figure S13). The nanotubes physical properties are shown in table 1.

**Figure 2.**
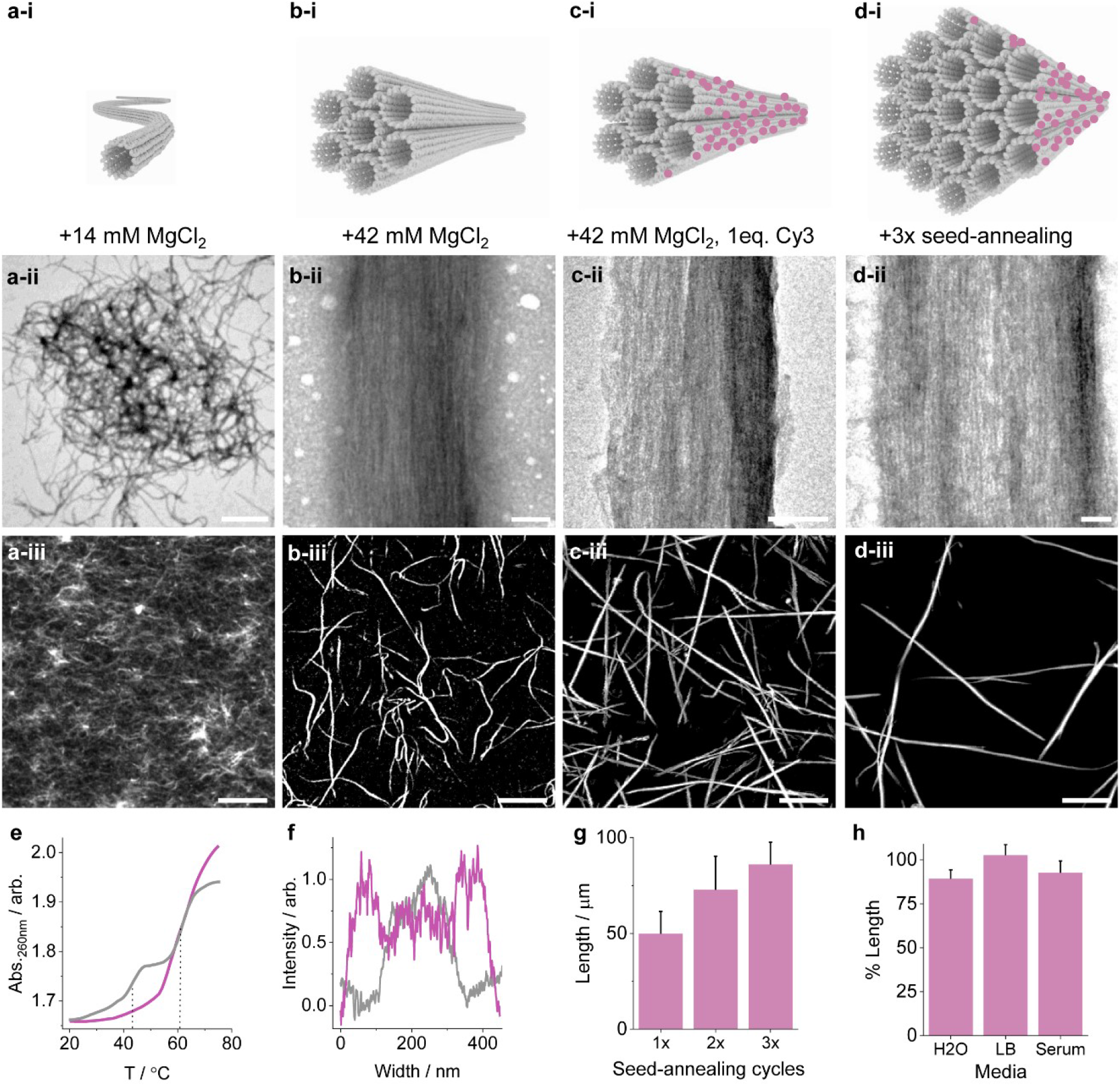
Structural characterization of DNA nanotubes and fibers. **a)** DNA nanotubes, **a-i)** 3D rendering, **a-ii)** TEM image of nanotube bundle, scale bar 1000 nm and **a-iii)** CLSM showing ensemble nanotube bundles, scale bar 10 μm. **b)** DNA fibers, **b-i)** 3D rendering, **b-ii)** TEM image of fiber, scale bar 50 nm and **b-iii)** CLSM image showing ensemble fibers, scale bar 10 μm. **c)** Ultra-rigid DNA fiber characterization using 1 eq. of Cy3 modification per subunit (magenta spheres), **c-i)** 3D rendering, **c-ii)** TEM image of fiber cross-section, scale bar 50 nm, **c-iii)** CLSM image showing ensemble fibers, scale bar 10 μm. **d)** Ultra-long DNA fiber characterization with multiple seed-annealing cycles containing 1 eq. of Cy3 modifications per subunit (magenta spheres), **d-i)** 3D rendering, **d-ii)** TEM image, scale bar 50 nm, **d-iii)** CLSM image of ensemble fibers, scale bar 10 μm. **e)** UV absorbance melting profiles at 260 nm comparing DNA nanotubes (grey line) and DNA fibers (magenta line), dotted lines represent the Tm values. **f)** TEM cross-section width analysis comparing DNA fibers without (grey line) and with 3x seed-annealing cycles (magenta line). **g)** CLSM-derived length analysis of DNA fibers with various seed-annealing cycles, values obtained by plotting the average longest fibers for each cycle. **h)** CLSM-derived stability of DNA fibers in deionized water (H2O), lysogney broth (LB) and fetal bovine serum (Serum), values represent the percentage length after 4 hr exposure at 37 °C.

**Table 1.** Physical properties of DNA nanotubes and fibers.

| Construct | Maximum length ($\mu\text{m}$) | Persistence length ($\mu\text{m}$) | $T_m$ (°C) |
| --- | --- | --- | --- |
| DNA nanotubes | - | $2.0 \pm 0.1$ | 47.1, 62.0 |
| DNA fiber 0.1 eq. Cy3 | $33.3 \pm 3.7$ | $17.9 \pm 1.7$ | 61.1 |
| DNA fiber 1 eq. Cy3 | $40.6 \pm 6.3$ | $217.2 \pm 31.4$ | 63.0 |
| 3x seed-anneal DNA fiber 1 eq. Cy3 | $101.1 \pm 29.8$ | $272.8 \pm 13.8$ | - |

At high divalent metal ion concentrations the DNA nanotubes coalesce to form highly rigid fibers (Figure 2b, Figure S1-3, S7-12). Structural characterization reveals long filaments are generated when the component mixture is annealed in at least 42 mM MgCl2 or CaCl2 (Figure S9). TEM analysis shows fibers are composed of tightly packed and aligned DNA nanotubes (Figure 2b-ii, Figure S13-14). The filaments measure approximately 7 nanotubes across, implying each fiber is a bundle of 37 condensed nanotubes. The nanotube packing arrangement is due to high concentrations of divalent metal cations which complex between adjacent phosphate anions from neighboring nanotubes (Figure 1a, inset). The divalent requirement is confirmed since monovalent NaCl_2_ and KCl_2_ do not form fibers (Figure S9). CLSM shows the fibers measure up to 28.1 μm ± 6.2 μm (Figure 2b-ii), and are almost an order of magnitude stiffer than isolated nanotubes (table 1, Figure S15-16). Agarose gel electrophoresis and a centrifugal pelleting assay confirms the fibers form in high ratios (Figure S7-8). Gel electrophoretic analysis shows the fibers do not migrate into the gel lane. This indicates the majority of the component oligonucleotides and subunits form a large molecular weight complex. A centrifugal pelleting assay backs up this finding which reveals 90.3 ± 2.3 % was large enough to precipitate under modest centrifugal forces.

The condensation strategy is compatible with other DNA nanotube designs. A tile^[16]^ and origami-based^[25]^ nanotube designs were assayed and their fiber-forming properties identified (Figure S17). The DNA tile is assembled from overlapping subunits, whilst the DNA origami structure contains terminal blunt-ends which interact with other subunits to form a nanotube array. CLSM shows both types of nanotubes are able to form fibers under high divalent metal ion concentrations, to confirm the condensation approach is generalizable.

The fibers’ length and stiffness can be improved by introducing high concentrations of aromatic fluorophores (Figure 2c). Cy3 is known to form homodimers *via* favorable π-stacking interactions.^[26]^ Increasing the modified DNA concentration from 0.1 μM to 1 μM, which equates to 0.1 and 1 Cy3 per subunit, respectively, increases the length and rigidity of fibers markedly. TEM analysis shows the fibers form tightly bound nanotube complexes with co-aligned subunits (Figure 2c-ii, S14). CLSM identified the longest fibers’ increased to over 40 μm, whilst the persistence length increased to over 200 μm (Figure 2b-ii, table 1, Figure S15-16), which is an order of magnitude stiffer than cytoskeletal actin filaments.^[27]^ The Cy3 dyes interact constructively in the nanotube bundle with adjacent dyes *via* π-stacking interactions. A similar stabilizing effect has also been observed for other covalently attached aromatic modifications on DNA, including porphyrins.^[28]^

Thermal UV absorbance and corresponding CLSM analysis confirms all DNA constructs remain structurally stable at physiological temperatures (Figure 2e, table 1, Figure S18). The DNA nanotubes and fibers were folded by thermally annealing between 75 °C to 20 °C at a rate of 1 °C per min. Monitoring the absorbance at 260 nm, the annealing profiles reveal a 2-step transition for all constructs. The higher annealing temperature (Ta) corresponds to the subunit formation, whilst the lower value corresponds to the nanotube or fiber formation step. This result implies both the nanotubes and fibers form simultaneously, and therefore follows a nucleation-elongation mechanism (Figure S18-19). Contrastingly, in the melting profiles, only DNA nanotubes display a 2-step melting transition (T_m_) (47.1 and 62.0 °C), whilst the fibers display single melting transitions (61.1-63.0 °C). The high thermal stability of the fibers indicates they form highly cooperative superstructures suitable for various bionanotechnology applications.

The fibers’ length and diameter can be improved by performing multiple seed-annealing cycles (Figure 2d, S20). The large hysteresis shift between the annealing temperature (~ 47 °C) and the melting temperature (~ 62 °C) can be exploited, enabling assembled fibers to be combined with additional component strands at 55 °C. Upon cooling, the supplemented oligonucleotides complex to the annealed fibers to increase their length and width markedly (Figure 2d-i-ii, S15, S20). Using this strategy, the fibers’ dimensions can be modulated by varying the number of seed-annealing cycles. 3x seed-annealing cycles improves the fibers’ stiffness further, with fibers over 100 μm long and 300 nm wide observed (table 1, Figure S13).

DNA fibers are stable in biocompatible cell media and display good nuclease resistance properties. The fibers were pelleted and resuspended in lysogney broth (LB) or fetal bovine serum (FBS), and the length monitored using CLSM (Figure 2g, S21-22). After 4 h, the length was measured to be >80 % of the original. This result confirms the fibers remain stable and resilient to physiological temperatures and proteins in both media types, including DNA digestion enzymes within this timeframe. The fibers also remain stable when resuspended into deionized water.

### 2.2 Functionalizing DNA fibers with synthetic biomolecules

DNA fibers can be modified with a range of biomolecules to introduce nanoscale functionality. Firstly, DNA single-strand linkers were incorporated to enable subsequent downstream DNA attachment (Figure 3a, see table S1-3 for sequences and mixing, and Figure S4 for 2D strand map). The extended linker strand contains the DNA sequence of a component oligonucleotide, but extended in the 5’ direction, leaving a single strand domain protruding from the fiber. TEM and CLSM shows the linker strands do not interfere with the fiber formation, even when added in a one equivalent stoichiometric ratio per subunit (Figure 3a-ii-iii, Figure S23-24). The linker regions can be used as aptamer sequences to direct cellular binding,^[29]^ or as DNAzymes to introduce subsequent mechanical action.^[30]^

**Figure 3.**
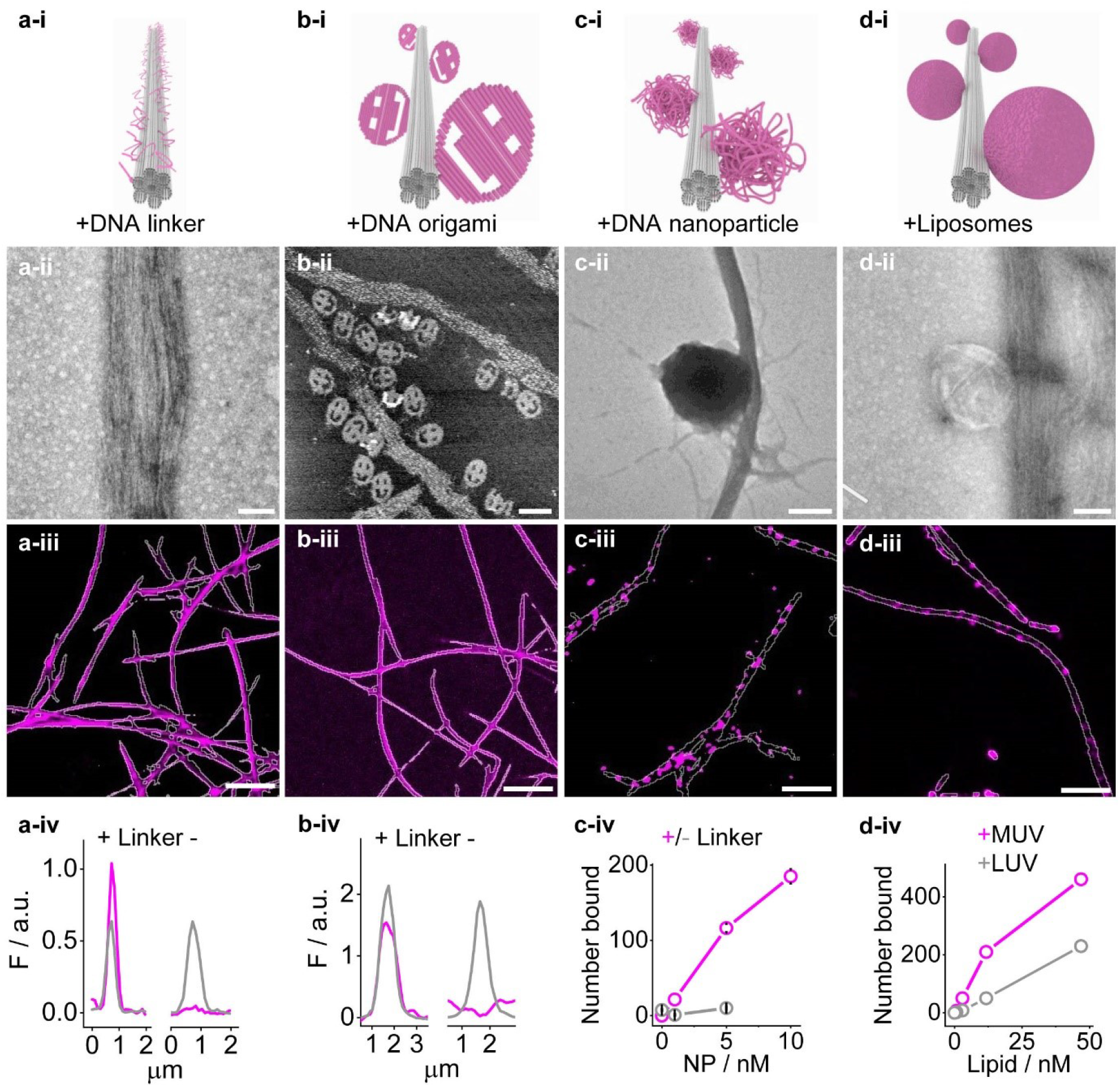
Functionalizing DNA fibers with synthetic biomolecules. **a)** Characterization of DNA fibers (grey) modified with linker strands (magenta) extended from component oligonucleotides, corresponding **a-i)** 3D rendering, **a-ii)** TEM image, scale bar 100 nm, **a-iii)** CLSM image of DNA fibers (grey channel) with fluorescein labelled-linker strands (magenta channel), scale bar 10 μm, and **a-iv)** cross-sectional analysis of CLSM images comparing DNA fibers (grey line) with and without linker strands (magenta line). **b)** Characterization of DNA fibers (grey) modified with fluorescein-labelled DNA origami smileys (magenta) connected *via* linker strands, corresponding **b-i)** 3D rendering, **b-ii)** AFM image of ruptured nanotubes, scale bar 100 nm, **b-iii)** CLSM image of DNA fibers (grey channel) with linker strands and DNA origami smileys (magenta channel), scale bar 10 μm, and **b-iv)** cross-sectional analysis of CLSM images comparing DNA fibers (grey line) with and without linker strands containing DNA origami smileys (magenta line). **c)** Characterization of DNA fibers (grey) modified with Cy5-labelled hydrophobic DNA nanoparticles (magenta) connected *via* linker strands, corresponding **c-i)** 3D rendering, **c-ii)** TEM image, scale bar 100 nm, **c-iii)** CLSM image of DNA fibers (grey channel) with linker strands and DNA nanoparticles (magenta channel), scale bar 20 μm, and **c-iv)** CLSM analysis comparing number of DNA nanoparticles bound along DNA fibers with (magenta line) and without linker strands (grey line). **d)** Characterization of DNA fibers (grey) complexed with vesicles (magenta) of different sizes, corresponding **d-i)** 3D rendering, **d-ii)** TEM image, scale bar 100 nm, **d-iii)** CLSM image of DNA fibers (grey channel) complexed with MUVs (magenta channel), scale bar 20 μm, and **d-iv)** CLSM analysis comparing the number of bound SUVs (grey line) vs MUVs (magenta line) along DNA fibers (grey line) with increasing concentration.

Secondly, DNA origami nanostructures can be attached to fibers *via* single strand linkers (Figure 3b). A dye-labelled DNA origami smiley measuring ~ 90 x 70 x 2 nm was incorporated into the DNA fibers by mixing the two folded and purified constructs (see figure S5 for 2D strand map, and table S4 for sequences, Figure S25). Atomic force microscopy (AFM) and CLSM confirmed the two synthetic biomolecules are able to assemble readily, and that the DNA origami is able to tether specifically to the fibers (Figure 3b-ii-iii, S26). A control fiber without the linker strands did not show any significant DNA origami loading, confirming the two complementary strands extended from the fiber and origami are able to form a stable duplex. This result means the DNA matrix can be decorated with a vast array of complex structures, from sensors,^[6]^ robots^[4]^ or quantum dot loaded motifs.^[20]^

Thirdly, hydrophobic DNA nanoparticles are able to bind along the DNA framework *via* single strand linkers (Figure 3c). DNA nanoparticles containing cholesterol-rich regions were generated from M13mp18 single strands (see Figure S6 for 2D strand map and S27, table S1-3 and S5 for sequences and mixing). Cholesterol-modified oligonucleotides were introduced to the scaffold by binding to 34 target sites using 17 helper strands. The embedded cholesterols interact with other functionalized scaffolds *via* favorable hydrophobic intermolecular interactions to form nanometer-sized particles. A single strand region of the M13mp18 scaffold was designed to tether to the extended linking strand from the fiber. TEM and CLSM showed the DNA nanoparticles are able to bind along the fiber *via* the linker strand (Figure 3c-ii-iii and S28-29). The role of the linker strand was confirmed since a control fiber without the linkers showed significantly less nanoparticle loading. The DNA nanoparticles may serve as sensitive functional sensors to support biomedical research along fibers.^[31]^

Finally, DNA fibers can be decorated with liposomes of various diameters (Figure 3d). Liposomes composed of lipid 1,2-dimyristoyl-sn-glycero-3-phosphocholine (DMPC) complex to DNA fibers *via* favorable ionic interactions. Vesicles ranging from 100 nm to 20 μm in diameter were screened and the fiber-loading capabilities determined. TEM and CLSM showed the liposomes complex readily to the fibers (Figure 3d and S30-31). Small, medium and large unilamellar vesicles (S-, M-, LUVs), with diameters of 100, 400 and 1000 nm, respectively, align along the 1D arrays. In contrast, giant unilamellar vesicles (GUVs), with diameters between 2-20 μm, are too large to align, and instead cause the fibers to wrap around the vesicles. The binding event is mediated *via* magnesium ions since a control assay using potassium ions did not show any vesicle loading. This result confirms magnesium ions coordinate between negatively charged phosphate anions from the DNA and negatively charged DMPC lipid headgroups. The immobilized liposomes can direct binding *via* lipid modified headgroups, or be used as active ingredient reservoirs for downstream cell-matrix applications.^[32]^

### 2.3 Aligning *E.coli* cells along DNA fibers

DNA fibers can be oriented under flow using microfluidic chambers. The fibers can be aligned on a glass surface by applying shear stress forces using a microfluidic flow slide. Time-series CLSM analysis shows fibers stick rapidly to the glass surface and align with the flow direction (see Figure S32). The aligned fibers average length is 57.2 (± 30.2) μm long.

Divalent metal cations mediate *E.coli* cell binding along flow-aligned fibers following a type I mechanism (Figure 1d). In nature, neutrophils excrete DNA to act as extracellular traps to capture Gram-negative pathogens in the presence of divalent metal ions^[33]^ Inspired by this process, magnesium and calcium ions were employed to mediate binding between negatively charged phosphates from the fiber, and negatively charged entities from the outer membrane of *E.coli*, including phosphatidylglycerol (PG) lipids and glycocalyx molecules (Figure 4a).^[34]^ TEM analysis shows the fibers are able to complex *E.coli* cells non-site-specifically along the 1D array in the presence of these cations (Figure 4b and S33). The images also reveal the fibers remain structurally rigid during the complexation process and can support micron-sized bacteria cells. CLSM was used to investigate the cation binding dependency of *E.coli* cells. Three salt conditions were screened to identify the optimal cell loading (Figure 4c, S34), divalent magnesium and calcium ions, and monovalent potassium ions. Using flow-aligned fibers, combined calcium and magnesium ions exhibit the most efficient *E.coli* cell complexation properties. Whilst potassium ions did not show any significant binding to confirm the divalent requirement. CLSM also revealed the cell binding event is rapid. Analysis of the combined entities in the presence of magnesium and calcium ions showed complexation occurred within seconds using a microfluidic flow chamber. Once bound, particle tracking analysis showed the cell remained fixed at the fiber site (Figure S35).

**Figure 4.**
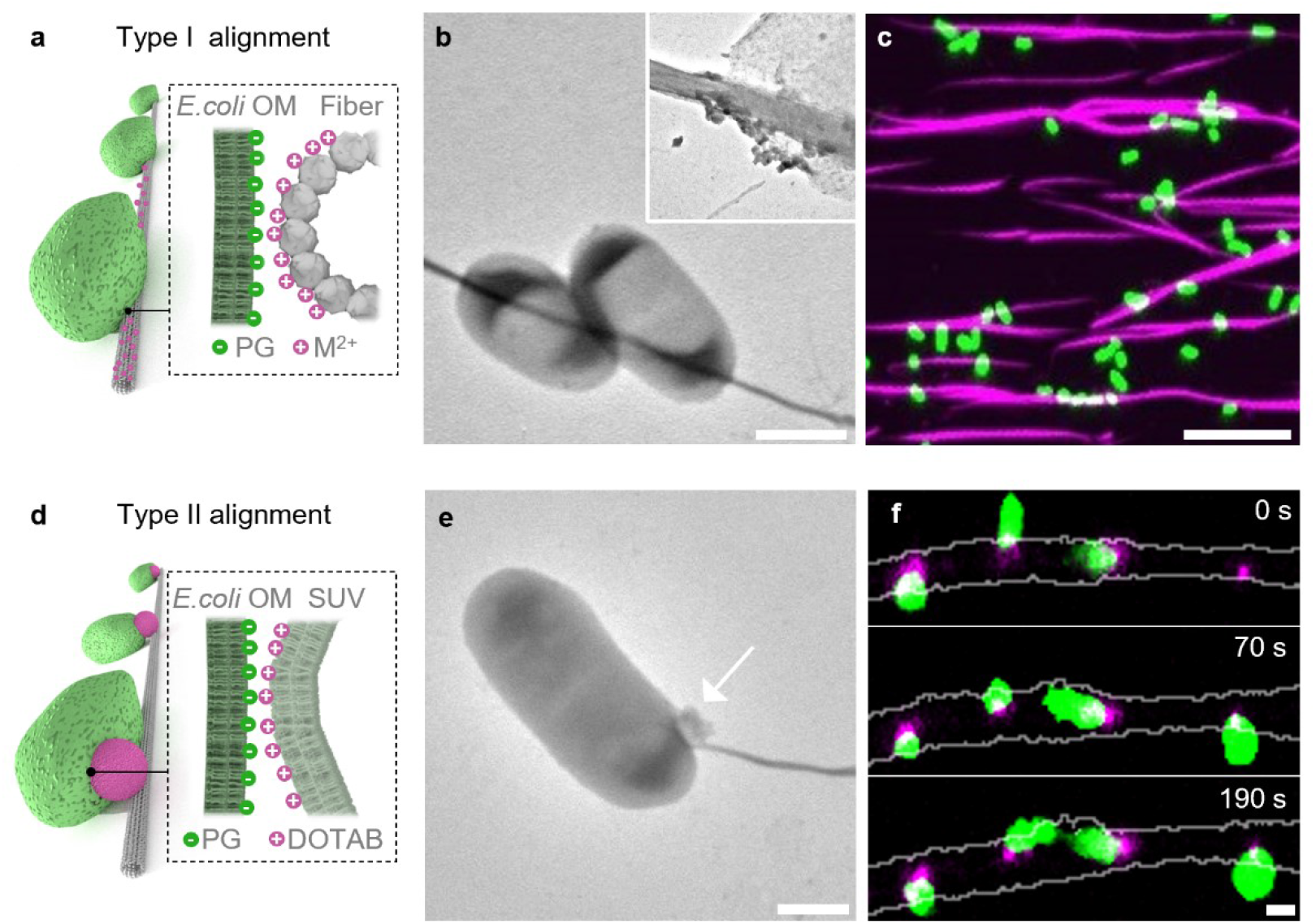
*E.coli* cells align along DNA fibers. Type I cell alignment mediated *via* Mg^2+^ and Ca^2+^ ions between DNA fibers (grey) and phosphatidylglycerol (PG) lipids found in the outer membrane (OM) of *E.coli* cells (green), corresponding **a)** 3D representation of complex, **b)** TEM image of fiber-*E.coli* complex in Mg^2+^ and Ca^2+^ solution, scale bar 1 μm, insert magnification of complex, **c)** CLSM image of flow-aligned fibers (magenta channel) complexed with *E.coli* cells (green channel) in Mg^2+^ and Ca^2+^ solution, scale bar 10 μm. Type II cell alignment mediated *via* cationic liposomes (magenta) between negatively charged DNA fibers (grey) and phosphatidylglycerol (PG) lipids found in the OM of *E.coli* cells (green), corresponding **d)** 3D rendering of complex, **e)** TEM image of *E.coli*-SUV-fiber, arrow denotes the position of the vesicle, scale bar 1 μm, **f)** CLSM time series of site specific loading of *E.coli* cells (green channel) at cationic SUV sites (magenta channel) tethered to DNA fibers (grey channel), scale bar 1 μm.

Cationic liposomes support type II *E.coli* cell alignment along DNA fibers. Positively charged lipids were introduced into liposomes^[35]^ to mediate binding between negatively charged DNA fibers and *E.coli* cells in a location-specific fashion (Figure 1d, 4d). Firstly, the optimal cationic-vesicle fiber loading ratio was identified using CLSM (see Figure S31). This was performed to ensure the vesicles remain isolated along the fibers. Next, a high concentration of bovine serum albumin (BSA) was added to the reaction mixture to minimize cellular binding *via* Mg^2+^ ions. The serum proteins also prevent fibers from sticking to the glass surface, which has the added benefit of improving their molecular accessibility to freely diffusing cells. TEM and CLSM images shows the *E.coli* cells bind only at the vesicle sites along the DNA fiber (Figure 4e-f, S36-37). Two different 1,2-dioleoyl-3-trimethylammonium-propane (DOTAP)-containing vesicle sizes were assayed, 100 nm SUVs and 400 nm MUVs. The analysis revealed the smaller vesicles were not suitable for location-specific cellular binding, as each cell can bind to more than one vesicle along the fiber. In contrast, the larger vesicle size showed improved binding properties, where each cell can adhere to one vesicle along the fiber. Time-series particle tracking analysis shows the bound cells remain tethered but can diffuse around the liposome (Figure S38). The rotational movement is most likely due to lipid diffusion from the vesicle and cell membranes.^[36]^ Using this approach, the intercellular spacing between the immobilized cells can be identified. Gap analysis reveals the cells fluctuate with low micron-scale mobility.

### 2.4 Biological applications of *E.coli-DNA* fibers

DNA fibers can function as advanced extracellular scaffolds. The fibers can act as bacteria-capture devices and are able to bind *E.coli* cells under complex biological conditions. To test the cellular selectively, fibers were added to *E.coli* cells mixed with dilute whole blood (WB) in reduced serum media. Using a microfluidic chamber, CLSM reveals DNA fibers adhere *E.coli* cells exclusively and not erythrocytes or white blood cells (Figure 5a, S39). This selectively is due to blood cells having significantly reduced levels of negatively charged lipid molecules compared to *E.coli*.^[34]^ In addition, the analysis confirms the binding is maintained in the presence of serum proteins. At high concentrations the fibers form 3D networks which can act as conventional extracellular matrices.^[37]^ CLSM shows the fibers form interconnected frameworks when left to settle on a surface at 1 μM or higher. When *E.coli* cells are added to Mg^2+^ and Ca^2+^ coated fibers, the cells stick readily *via* a type I mechanism (Figure 5b, S40). Importantly, the bound cells are viable for both alignment systems. Times series analysis shows the vesicle-fused cells are viable (Figure S41). After 29 min of cell adhesion, the images confirm the *E.coli* cells are able to divide at the vesicle site. Contrastingly, for calcium-mediated alignment, the dividing cells can remain complexed to the extracellular scaffold. Using non-flow aligned fibers, CLSM shows the cells are able to divide sequentially along the DNA framework (Figure 5c, S42). The image confirms the 1D array can sculpt the dividing daughter cells to enable long-term multi-generational order.

**Figure 5.**
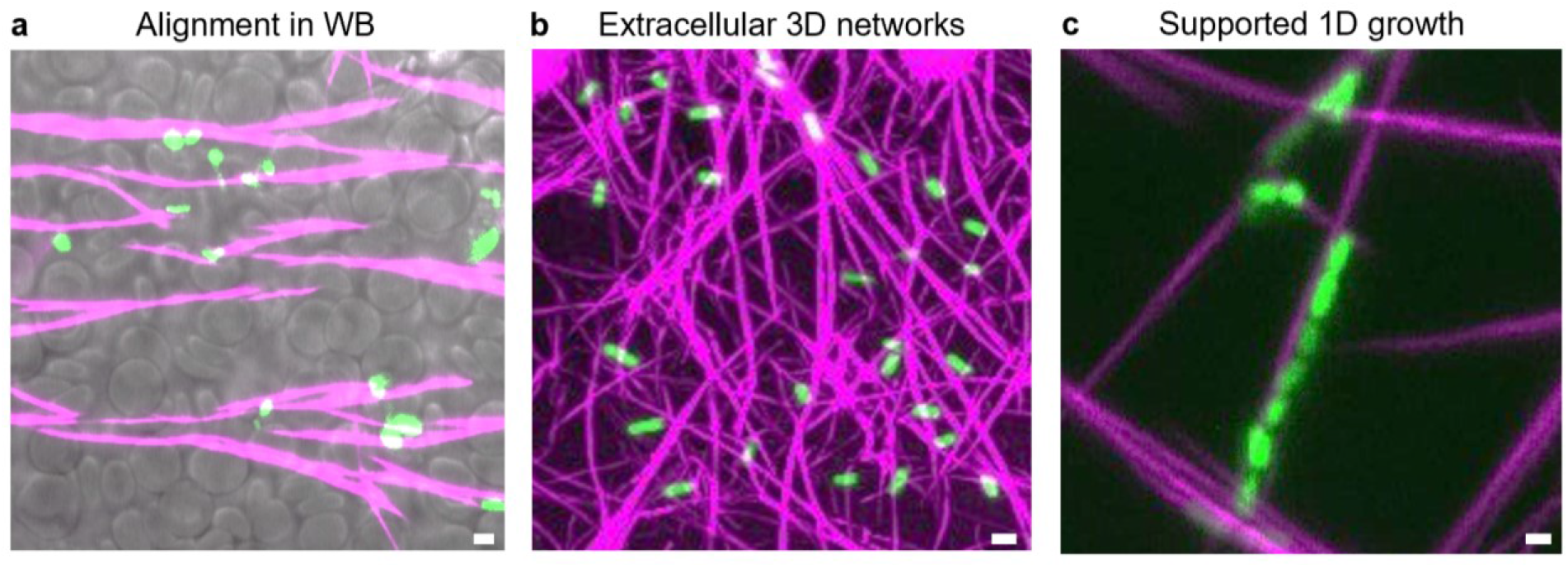
Applications of extracellular DNA fibers. **a)** DNA fibers (magenta channel) selectively adhere *E.coli* cells (green channel) and not erythrocytes or white blood cells found in whole blood (WB, bright-field channel) in reduced serum *via* a type I mechanism using Ca^2+^ and Mg^2+^ ions, scale bar 2 μm. **b)** 3D DNA fiber matrix (magenta channel) aligning *E.coli* cells *via* a type I mechanism using Ca^2+^ and Mg^2+^ ions, scale bar 2 μm. **c)** Over time DNA fibers support multi-generational alignment of dividing *E.coli* cells, scale bar 2 μm.

## 3.0 Summary and perspective

A new macromolecular nanoarchitectonic approach has been developed which can align synthetic biomolecules and *E.coli* cells. The findings expand the boundaries of DNA nanotechnology by generating highly functional macroscopic fibers. The results show DNA nanotube condensation is very efficient, robust and versatile. The strategy is generalizable as both DNA origami and tile-based nanotube designs can be condensed into fibers. Modifying the surface chemistry of the fibers by introducing π-stacking modifications, or adapting the annealing protocol, enables the diameter, length and rigidity to be customized. However, there is scope to expand the properties further, for instance, applying more seed-annealing cycles should yield even longer and stiffer fibers. Alternative condensation agents, like diverse divalent metals or small cationic organic molecules^[24]^ can enhance the fibers chemical diversity. DNA fibers may be simplified by using nanotubes comprised of just one oligonucleotide.^[15]^ Or contrastingly, employing origami-based designs, which exploit secondary structure-recognition^[9]^, may facilitate sequence-specific control across the entire fiber length.

The biophysical ability to organize bacteria cells along DNA fibers should find widespread use in biomedical research. The results show DNA fibers’ capture selectivity is maintained in complex biological systems paving the way for future clinical applications, including, novel wound-dressing materials to dialysis devices.^[38]^ The intercellular-gap analysis between tethered *E.coli* cells reveals their position is relatively fixed along the framework. This ability means the proximity between bound cells can be determined and important studies performed, such as, distance thresholds for cell-cell communication, the generation of complex 3D cell matrixes during biofilm formation, to aligned multi-generational cell experiments to study antimicrobial resistance.^[39]^ Incorporating advanced synthetic biomolecules along the DNA framework can help support mechanistic research, or be used to direct cell-specific binding. Smart molecules readily incorporated include, DNA aptamers, DNA origami-based robots or liposome agents, all of which can be activated to release payloads on command. In addition, the extracellular scaffold can be selectively digested by applying a high concentration of DNAse enzymes. These advanced matrix properties are not readily possible with existing extracellular scaffolds.

In synthetic biology, new functional materials can be generated, including replicating actin filaments inside model cells with controlled cellular movement, to mimicking flagella motion of synthetic bacteria. Alternatively, with bigger and stiffer fibers, the arrays should be able to support mammalian cells, opening the route for unique cell-, skin- and organ-grafting materials.^[40]^ Importantly, the unit cost of the DNA framework is relatively cheap, requiring just 5 component single strands, meaning the new materials can be applied to many approaches at scale. To complement this, the fibers can be pelleted readily using centrifugation, and therefore concentrated, diluted or exchanged into new buffer systems in seconds. In conclusion, my report advances the field of macromolecular nanoarchitectonics across three disciplines, and helps to address the demand for macroscopic materials with nanotechnology-enabled functionality.

## Acknowledgements

Prof. Stefan Howorka for kindly providing laboratory equipment and support. Prof. Mervyn Singer and Dr. Nishkantha Arulkumaran for providing human whole blood under license.

## Additional Information

Supporting information and supplementary data is available online.

